# Histologic safety of transcranial focused ultrasound neuromodulation and magnetic resonance acoustic radiation force imaging in rhesus macaques and sheep

**DOI:** 10.1101/827063

**Authors:** Pooja Gaur, Kerriann M. Casey, Jan Kubanek, Ningrui Li, Morteza Mohammadjavadi, Yamil Saenz, Gary H. Glover, Donna M. Bouley, Kim Butts Pauly

## Abstract

**Background:** Neuromodulation by transcranial focused ultrasound (FUS) offers the potential to non-invasively treat specific brain regions, with treatment location verified by magnetic resonance acoustic radiation force imaging (MR-ARFI).

**Objective:** To investigate the safety of these methods prior to widespread clinical use, we report histologic findings in two large animal models following FUS neuromodulation and MR-ARFI.

**Methods:** Two rhesus macaques and thirteen Dorset sheep were studied. FUS neuromodulation was targeted to the primary visual cortex in rhesus macaques and to subcortical locations, verified by MR-ARFI, in eleven sheep. Both rhesus macaques and five sheep received a single FUS session, whereas six sheep received repeated sessions three to six days apart. The remaining two control sheep did not receive ultrasound but otherwise underwent the same anesthetic and MRI procedures as the eleven experimental sheep. Hematoxylin and eosin-stained sections of brain tissue (harvested zero to eleven days following FUS) were evaluated for tissue damage at FUS and control locations as well as tissue within the path of the FUS beam. TUNEL staining was used to evaluate for the presence of apoptosis in sheep receiving high dose FUS.

**Results:** No FUS-related pre-mortem histologic findings were observed in the rhesus macaques or in any of the examined sheep. Extravascular red blood cells (RBCs) were present within the meninges of all sheep, regardless of treatment group. Similarly, small aggregates of perivascular RBCs were rarely noted in non-target regions of neural parenchyma of FUS-treated (8/11) and untreated (2/2) sheep. However, no concurrent histologic abnormalities were observed, consistent with RBC extravasation occurring as post-mortem artifact following brain extraction. Sheep within the high dose FUS group were TUNEL-negative at the targeted site of FUS.

**Conclusions:** The absence of FUS-related histologic findings suggests that the neuromodulation and MR-ARFI protocols evaluated do not cause tissue damage.

## Introduction

Transcranial focused ultrasound (FUS) delivers targeted ultrasound energy to specific brain regions without damaging intervening tissue or requiring skull removal (Martin and Werner, 2013; Lipsman et al., 2014). Importantly, transcranial FUS avoids the risks associated with invasive procedures (*e.g.*, bleeding, infection) while maintaining high spatial resolution and the ability to reach subcortical targets, which limit other neurosurgical and neurostimulatory methods.

A potentially transformative application of transcranial FUS is neuromodulation, which is thought to be a noninvasive method to explore brain function and circuitry (Tyler et al., 2018). Neuromodulation uses short bursts of low intensity ultrasound to excite or inhibit neural activity and can be targeted to subcortical structures at the scale of a few millimeters, which cannot be achieved by other noninvasive neuromodulation modalities such as transcranial magnetic or electrical stimulation (Monti et al., 2016; Naor et al., 2016; Fomenko et al., 2018). This could enable functional mapping of small nuclei for treatment targeting and for advancing neuroscience, and offer a possible treatment for neurological conditions (Kubanek, 2018).

Human studies of FUS neuromodulation of cortical and subcortical regions have not led to detectable tissue changes on post-session MRI or behavioral deficits (Hameroff et al., 2013; Lee et al., 2015, 2016b; Legon et al., 2014, 2018a,b). As summarized in a recent review of FUS neuromodulation, fourteen out of fifteen animal publications showed no abnormal histologic findings (Blackmore et al., 2019). Included in the fourteen studies were two large animal studies, one in pigs (Dallapiazza et al., 2017) and one in macaques (Verhagen et al., 2019), which found no tissue damage resulting from FUS neuromodulation. However, one study in sheep raised concerns of microhemorrhage after exposure to prolonged, repetitive FUS neuromodulation (Lee et al., 2016c). Thus, the first purpose of this work was to ascertain whether neuromodulation poses a risk of tissue microhemorrhage in sheep as suggested by Lee *et al.*, with the addition of controls not treated with FUS, and in rhesus macaques.

In addition, FUS neuromodulation is aided by confirmation of FUS targeting in the brain. MR acoustic radiation force imaging (MR-ARFI) uses a series of very short FUS bursts at higher intensity to visualize the ultrasound focal spot *in situ*. The ultrasound pulses slightly displace tissue which, in synchrony with MRI, can be detected as a shift in image phase (McDannold and Maier, 2008). This phase shift is proportional to the ultrasound intensity applied, and therefore can provide a non-invasive metric of the intensity delivered at the focal spot. MR-ARFI can also be used to assess and compensate for distortion of the ultrasound through the skull. Proposed clinical applications of MR-ARFI include validation of treatment targeting (Holbrook et al., 2011; Auboiroux et al., 2012), optimization of transducer focusing through the skull (Larrat et al., 2009; Marsac et al., 2012; Vyas et al., 2014), and assessment of tissue changes during treatment (McDannold and Maier, 2008; Holbrook et al., 2010; Bitton et al., 2012).

Almost no assessments of MR-ARFI safety have been reported. Two reports of *in vivo* MR-ARFI in the body, one in rabbits (Huang et al., 2009) and one in pigs (Holbrook et al., 2011), have been published but did not discuss safety. One study involving transcranial MR-ARFI in two macaques has been published, but did not include histology (Chaplin et al., 2019). To our knowledge, the only report of MR-ARFI safety is from a study that investigated histology after transcranial MR-ARFI in one rodent, in which no tissue damage was observed (Larrat et al., 2009). The second purpose of this work was to assess tissue safety in a controlled study of transcranial MR-ARFI in sheep.

We evaluate histology in brain tissue following FUS neuromodulation in the visual cortex of rhesus macaques, and following neuromodulation and MR-ARFI in subcortical brain regions in sheep. The sheep histology includes a treatment control group in which no FUS was applied, and internal controls from hemispheres not treated with FUS. Our neuromodulation protocols included a component similar to those used in human studies, and to those evaluated by Lee and colleagues. We also investigated a broader range of intensity values and repeated number of FUS bursts, exceeding those values typically used in human protocols as well as those used in the study by Lee *et al*. Our findings provide important information for subsequent studies involving FUS neuromodulation or MR-ARFI.

## Materials and Methods

All animal experiments were performed with institutional approval from the Stanford University Administrative Panel on Laboratory Animal Care.

### Rhesus macaque study

Two 4-year-old adult male rhesus macaques (4.6 kg and 4.8 kg) were acquired from the Wisconsin National Primate Research Center in November 2016. Both non-human primates (NHP-1 and NHP-2) were clinically healthy on physical examination and were seronegative for the following pathogens: *Mycobacterium tuberculosis*, simian immunodeficiency virus, and simian T-lymphotrophic virus type 1 and 2. One animal was seropositive for simian retrovirus. Animals were housed in indoor caging and maintained on a 12:12 hr light:dark cycle in an AAALAC-accredited facility. Animals were fed a commercial primate diet (Teklad Global 20% Protein Primate Diet 2050, Envigo, Madison, WI) supplemented with fresh produce, and had unrestricted access to water. Figure 1(a) summarizes study characteristics.

**Figure 1:**
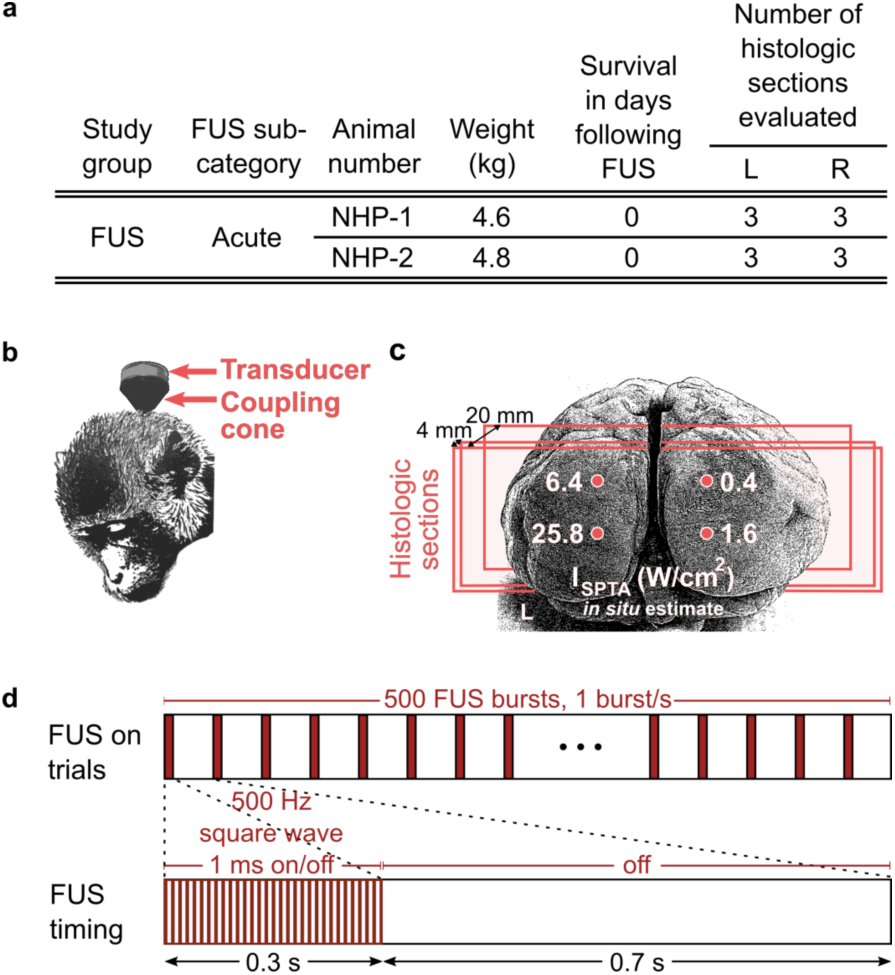
Summary of rhesus macaque study parameters. (a) Inclusion characteristics, survival time, and number of histologic samples evaluated for left (L) and right (R) hemispheres. (b) Illustration of rhesus macaque transducer positioning and (c) grid of focused ultrasound sonication in the visual cortex, where each location corresponds to estimated *in situ* spatial peak-temporal average intensity (I_*SPTA*_) values of 0.4, 1.6, 6.4, and 25.8 W/cm^2^, applied in short bursts. Vertical spacing between FUS targets was 10 mm (NHP-1) and 15 mm (NHP-2), and horizontal spacing was 15 mm (NHP-1) and 20 mm (NHP-2). The lower two target locations (1.6 and 25.8 W/cm^2^ I_*SPTA*_) were placed 2 mm above the inion. Three coronal histologic sections were obtained from each hemisphere of the visual cortex (approximate locations shown by red planes). The first histology plane was located near the cortical surface, the second at a depth of approximately 4 mm, and the third at a depth of approximately 20 mm. (d) Illustration of neuromodulation protocol comprising 500 FUS bursts.

#### Anesthesia and preparation

Both animals were sedated with ketamine (4 mg/kg, intramuscularly) and dexmedetomidine (0.02 mg/kg, intramuscularly) and anesthetized with 2-3% isoflurane throughout the FUS procedure. The hair was shaved from the back of the head prior to transducer placement.

#### Focused ultrasound

A single-element, 270 kHz focused ultrasound transducer fitted with an agar-filled cone was positioned at the back of the head and coupled with degassed ultrasound gel as illustrated in Fig. 1(b) (H-115, Sonic Concepts, Bothell, WA).

FUS was targeted to four regions in the visual cortex as shown in Fig. 1(c). A coupling cone was used such that the ultrasound focus was positioned at the surface of the brain (5 cm length from transducer). The focal pressure half-width was approximately 17 mm in the axial direction and 6 mm in the lateral direction. The lower two focal spot locations were placed 2 mm above the center of the inion and spaced bilaterally by 15 mm (NHP-1) and 20 mm (NHP-2). The upper two focal spot locations were located directly above at 10 mm (NHP-1) or 15 mm (NHP-2).

FUS was applied in 300 ms pulsed (50% duty cycle) bursts occurring every 1 s for a total of 500 stimuli, as illustrated in Fig. 1(d). One 8.3 min FUS trial (comprising 500 FUS bursts) was applied to each of the four neuromodulation locations. Free-field stimulus pressure levels corresponded to 0.5, 1, 2, and 4 MPa as measured in a water tank by fiberoptic hydrophone (Precision Acoustics, Dorset, UK), in order to sample a range of values. *In situ* intensity was estimated after assuming approximately 40% pressure loss through the macaque skull, based on reports from a previous study (Deffieux et al., 2013). One spatial peak-temporal average intensity (I_*SPTA*_) level was applied per location, with estimated *in situ* values of 0.4 (top) and 1.6 (bottom) W/cm^2^ on the right hemisphere and 6.4 (top) and 25.8 (bottom) W/cm^2^ on the left hemisphere, as illustrated in Fig. 1(c).

#### Fixation and histopathology

Thirty minutes following FUS, the animals were anesthetized to a surgical plane with 5% isoflurane and initially perfused with 0.25-0.5 liters of saline. Next, the macaques were perfused with 4 liters of 3.5% to 4% paraformaldehyde in 0.1 M phosphate buffer at high pressure for 2-3 minutes (2 liters) and at low pressure (2 liters) for one hour. Lastly, they were perfused with 1-1.25 liters each of 10%, 20%, and 30% sucrose solutions at high pressure for cryoprotection. The skull was removed using an autopsy saw (Shandon, ThermoFisher Scientific, No. 10000) and the brain was extracted. The primary visual cortex was segmented from the remaining cortex by making a coronal cut 2 mm posterior to the lunate sulcus. Brains were then immersion-fixed in 10% neutral buffered formalin for 7-10 days. Formalin-fixed tissues were then processed routinely, embedded in paraffin, sectioned at 7 *µ*m, and stained with hematoxylin and eosin (H&E). Three coronal tissue sections were obtained from each hemisphere of the visual cortex, resulting in six total sections per macaque (Fig. 1(c)). Each pair of left and right sections captured a crosssection of all four focal spot beams and covered the full extent of each hemisphere. The first two section pairs were obtained near the surface of the brain, in the region of the focal peak, spaced about 4 mm apart. The third section pair was located about 3 mm beyond the half-max intensity of the focus, at an approximate depth of 2 cm from the cortical surface. Slides were blindly reviewed by a board-certified veterinary pathologist (DB) for the presence of necrosis, apoptosis, edema, hemorrhage, inflammation, and neuropil rarefaction.

### Sheep study

Thirteen male Dorset sheep weighing 22 to 36 kg were included in the study. Eleven underwent transcranial FUS. Two animals did not receive ultrasound but otherwise under-went the same experimental procedures.

Sheep were divided into FUS (n=11) and control (n=2) study groups. Animals that received FUS were subdivided into four groups as follows: acute (n=2; euthanized zero days after FUS study), delayed (n=3; euthanized four to seven days after FUS study), repeated (n=3; underwent FUS again three to six days after the first FUS session, and euthanized four days after the last FUS study), and high dose (n=3; received multiple FUS sessions with prolonged application of FUS on the last day of study, and euthanized four days later). Both sheep in the control group underwent multiple days of MRI study. The two sheep in the acute FUS group and one sheep in the delayed FUS group also underwent MRI study on one or more days prior to the FUS session. Study characteristics are summarized in Fig. 2(a).

**Figure 2:**
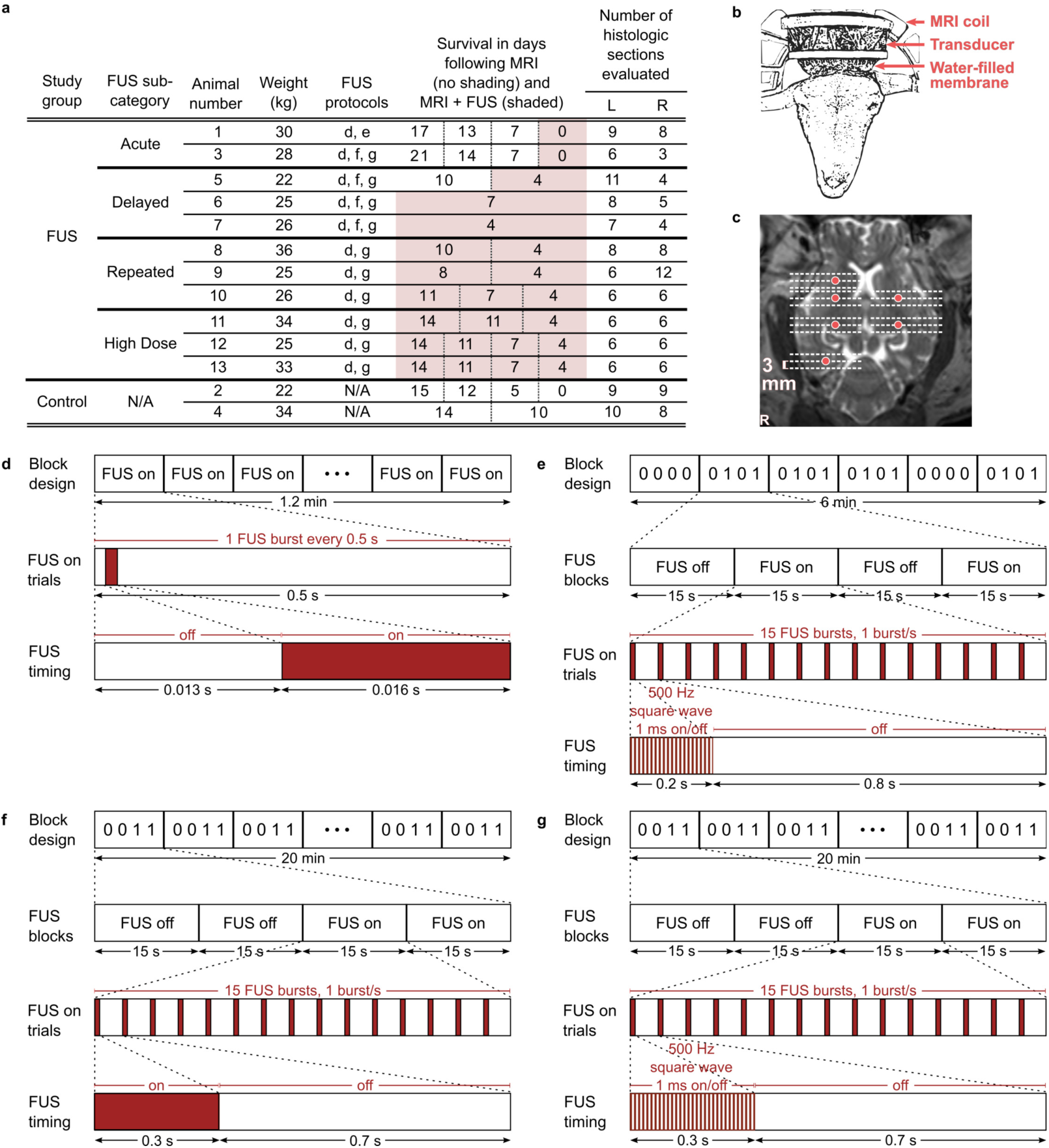
Summary of sheep study parameters. (a) Sheep inclusion characteristics. The two sheep in the control group underwent MRI and anesthesia but no FUS. The eleven sheep that underwent FUS were subdivided into acute (euthanized zero days after FUS study), delayed (euthanized four to seven days after FUS study), repeated (underwent multiple FUS sessions, and euthanized four days after the last FUS study), and high dose groups (underwent prolonged MR-ARFI applications at one location on the last day of study). Days of survival following the first (left-most) and subsequent days of study are reported in split columns where applicable, for MRI without FUS (unshaded cells) and MRI with FUS sessions (shaded cells). The number of evaluated histologic sections is directly related to the number of FUS targets per sheep. (b) Sheep transducer positioning and (c) exemplary focused ultrasound sonication locations (6 locations shown; red circles) shown on axial T2-weighted MRI (cropped to show detail). Histologic sections were obtained from each location targeted with focused ultrasound and additionally from planes approximately 3 mm rostral and caudal to targeted locations (18 sections shown; dashed lines). Illustration of (d) MR-ARFI focal spot localization and (e-g) neuromodulation FUS protocols. Protocols comprised (d) 128, (e) 120 and (f-g) 600 FUS bursts.

#### Anesthesia and preparation

Sheep were fasted for 24 hours prior to the study and then sedated with tiletamine and zolazepam (Telazol, Lederele Parenterals, Carolina, Puerto Rico) at 4 mg/kg, intramuscularly. Anesthesia was induced with a combination of 3% isoflurane in oxygen delivered by facemask and telazol in a continuous rate of infusion. All animals were orotracheally intubated and anesthesia was maintained with 1% to 3% isoflurane in oxygen with MRI conditional mechanical ventilation (Omni-Vent Series D, Allied Healthcare Products, St. Louis, MO) to maintain end-tidal carbon dioxide between 35 mm Hg and 45 mm Hg. Stomach tubes were placed after intubation to resolve gaseous distension and prevent regurgitation. Venous and arterial catheters were placed percutaneously for drug and fluid administration and blood pressure monitoring. Lactated Ringer’s solution (Abbott Laboratories, Abbott Park, IL) was administered intravenously at approximately 10 mL/kg/hr throughout anesthesia. The top of the head was shaved and treated with a depilatory cream for hair removal.

#### Physiological monitoring

Serial samples of hematocrit and arterial blood gases were taken from the auricular arterial catheter. Blood gas samples were analyzed immediately on a calibrated blood gas analyzer (i-STAT, Abbott Point of Care, East Windsor, NJ). Pulse oximetry measurements and capnography were performed continuously during anesthesia (Expression MR400, Philips Healthcare, Vantaa, Finland).

#### MR-guided focused ultrasound

MR-guided focused ultrasound studies were conducted using a 1024 element, 550 kHz focused ultrasound transducer fitted with a membrane containing chilled, degassed water (ExAblate 2100, Insightec Ltd., Haifa, Israel). The transducer was positioned above the head with degassed ultrasound gel for acoustic coupling (Fig. 2(b)).

Acoustic coupling and focal spot location were verified by MR-ARFI in the eleven sheep that underwent transcranial FUS. Figure 2(d) illustrates the MR-ARFI protocol in which FUS was on for 16 ms bursts within a 500 ms window (corresponding to the MR repetition time) over a period of 1.2 min. Each application of MR-ARFI comprised 128 FUS bursts. Figure 2(e-g) illustrates neuromodulation protocols, in which FUS was on for 200-300 ms bursts every 1 s with continuous wave (Fig. 2(f)) or pulsed (50% duty cycle) ultrasound (Fig. 2(e,g)). Each neuromodulation application comprised 120 (Fig. 2(e)) or 600 FUS bursts (Fig. 2(f,g)) over a period of 6 (Fig. 2(e)) or 20 minutes (Fig. 2(f,g)). The protocols applied for each sheep are reported in Fig. 2(a). FUS pulse timing was controlled by Eprime scripts (Psychology Software Tools, Pittsburgh, PA).

Multiple MR-ARFI and neuromodulation trials were administered consecutively to investigate the safety of repeated FUS sonications. The within-session timing of FUS application is illustrated in Fig. 3 for each sheep. Applied acoustic powers ranged from 127.5-195.5 W for MR-ARFI and 2-34 W for neuromodulation, and are summarized in Fig. 4(a) and Fig. 4(d), respectively, for each sheep. Neuromodulation acoustic powers were selected to result in at least 5.7 W/cm^2^ I_*SPTA*_ *in situ*, to replicate acoustic intensities applied in a study which reported tissue damage in sheep (Lee et al., 2016c), but to also include a broader intensity range to evaluate potential effects at higher levels.

**Figure 3:**
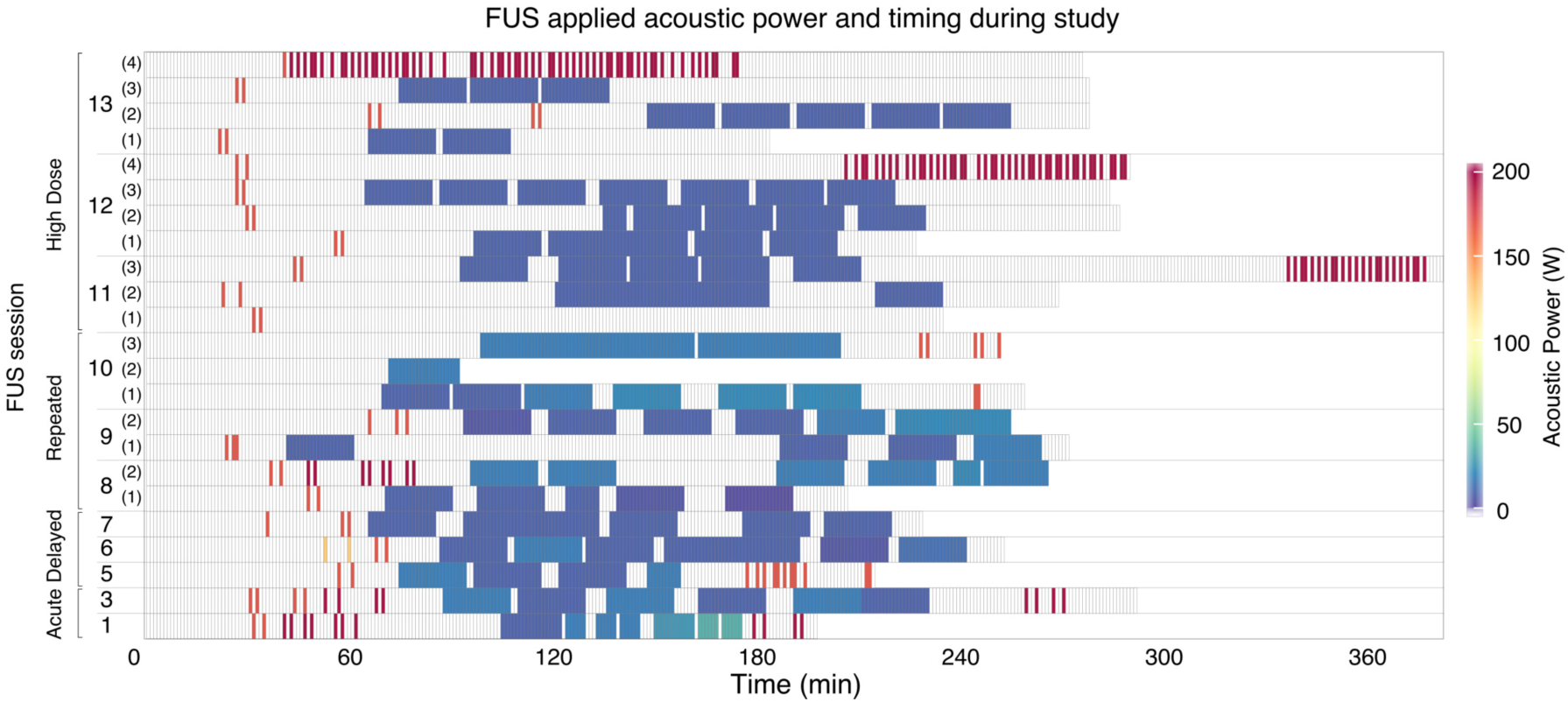
*In vivo* sheep study parameters. FUS applied acoustic power over time for each animal. Timing spans the total MRI and FUS session. Each cell represents a one minute interval, with color coding to indicate non-zero FUS acoustic powers. Empty cells indicate no FUS.

**Figure 4:**
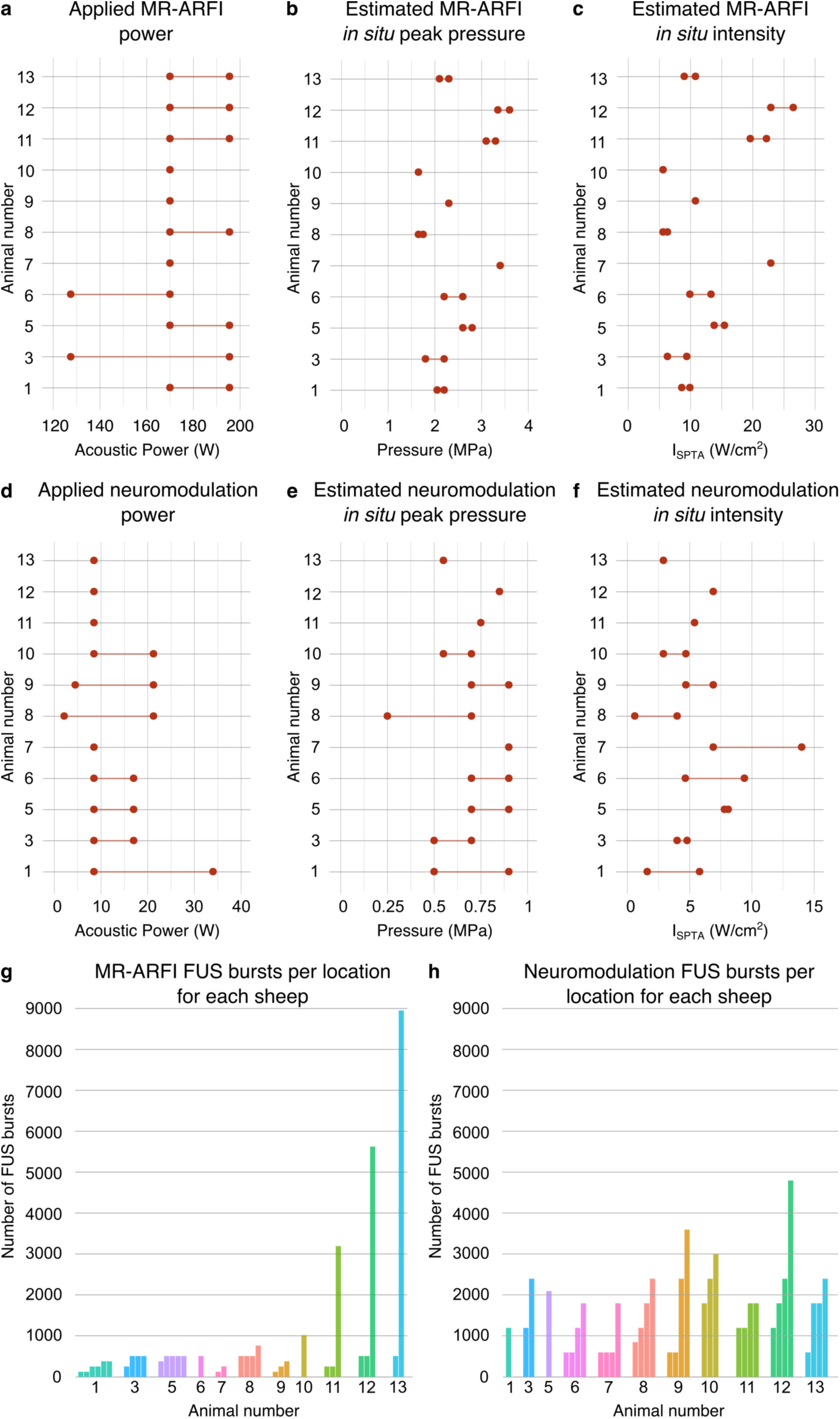
*In vivo* sheep study parameters. (a,d) Range of applied acoustic powers and estimated *in situ* (b,e) peak pressure and (c,f) spatial peak temporal average intensity for MR-ARFI and neuromodulation, respectively. Total number of FUS bursts applied to each (g) MR-ARFI and (h) neuromodulation location, where animal number is reported below each bar cluster. Individual bars represent unique sonication locations, and bar height indicates number of FUS bursts delivered to that location.

MR-ARFI and neuromodulation were targeted to 1-6 and 1-4 subcortical locations, respectively. The neuromodulation study measured visual evoked potentials using scalp electrodes in response to external stimulation (flashing lights) as well as during focused ultrasound sonication targeted to the visual pathway (lateral geniculate nucleus), the results of which are presented elsewhere (Mohammadjavadi et al., 2019). The lateral geniculate nucleus was a common neuromodulation location for all sheep, with additional focal spots typically located in planes approximately 10, 15, and 20 mm rostral and 10 mm caudal to the lateral geniculate nucleus. The focal pressure half-width was approximately 20 mm in the axial direction and 3.5 mm in the lateral direction. Figure 2(c) shows an example of targeted focal spot locations (sheep 9). The total number of FUS bursts applied to each targeted location are illustrated for MR-ARFI in Fig. 4(g) and for neuromodulation in Fig. 4(h), for each sheep. For the sheep in the repeated and high dose FUS groups, FUS locations were revisited for MR-ARFI and neuromodulation on multiple days. Two sheep had locations that were targeted both for MR-ARFI and neuromodulation on alternate days (two locations for sheep 8 and one location for sheep 9). Additionally, the three sheep in the high dose group each had one location that received MR-ARFI and neuromodulation during the same session. At the conclusion of the study, a high number of consecutive MR-ARFI repetitions were targeted to a single location in the high dose group, bringing the total number of MR-ARFI repetitions to 25, 44, and 70 at a single location (sheep 11, 12, and 13, respectively). Target locations were in the left hemisphere for acute and delayed groups, and bilateral for the repeated and high dose FUS groups.

#### MR imaging

MR-guided focused ultrasound studies were performed at 3T (Signa Excite, GE Health-care, Milwaukee, WI) using a quadrature head coil. A high resolution T2-weighted sequence was acquired for treatment planning with 2.5 s repetition time, 72 ms echo time, 22 cm isotropic field of view, and 256×192 acquisition matrix. MR-ARFI was performed using a spin echo sequence with repeated bipolar motion encoding gradients, 2DFT read-out, 500 ms repetition time, 39 ms echo time, 20×20×0.7 cm^3^ field of view, and 256×128 acquisition matrix (Bitton et al., 2012). Focused ultrasound application spanned from the second lobe of the first bipolar through the first lobe of the second bipolar motion encoding gradient. Images of the focal spot encoded by MR-ARFI were calculated by complex phase difference of two acquisitions with alternating motion encoding gradient polarities.

#### Histopathology and TUNEL

Animals were euthanized with a barbiturate overdose of 1 ml per 10 pounds of body weight of euthanasia solution (390 mg/mL pentobarbital and 50 mg/kg phenytoin, Virbac, St Louis, MO). Cardiac arrest was confirmed by auscultation. Skulls were removed via an autopsy saw (Shandon, ThermoFisher Scientific, No. 10000) and brains were extracted and immersion-fixed in 10% neutral buffered formalin for at least 10 days. Following fixation, the entirety of the brain was sectioned at approximately 3 mm intervals in the coronal plane. Brain regions were selected for histologic evaluation based on gross tissue comparison to MRI locations of FUS targets. Coronal tissue sections included the FUS target and all tissue dorsal to this region (to evaluate for potential cortical effects from skull heating and any effects within the FUS beam path). Additional tissue sections at distances of +/-3 mm from FUS targets were evaluated histologically (Fig. 2(c)). Tissue sections were also evaluated from contralateral, untreated hemispheres of acute and delayed FUS groups (internal controls). In control sheep, tissue sections were taken from the left and right hemispheres in locations anatomically similar to the FUS group. Formalin-fixed tissues were processed routinely, embedded in paraffin, sectioned at 5 *µ*m, and stained with H&E. Slides were blindly reviewed by a board-certified veterinary pathologist (KMC). Particular attention was paid to the presence or absence of hemorrhage, as well as pre-mortem tissue responses to damage (*i.e.*, necrosis, red blood cell engulfment (erythrophagocytosis), and intracellular red blood cell breakdown (hemosiderin-laden macrophages)). Additionally, terminal deoxynucleotidyl transferase-mediated dUTP-biotin nick end labeling (TUNEL) staining (ApopTag kit; Millipore, Temecula, CA) was performed according to manufacturer’s instruction on tissue sections corresponding to locations receiving the highest number of MR-ARFI repetitions from sheep in the high dose group.

#### Hydrophone measurements

*Ex vivo* skull caps from each sheep were degassed and placed in front of the focused ultrasound transducer array in a tank with degassed water. A fiberoptic hydrophone was positioned at the ultrasound focus to measure peak negative pressure transmitted through each skull cap to obtain an *in situ* intensity estimate for each acoustic power level applied *in vivo* (Precision Acoustics, Dorset, UK).

## Results

### Rhesus macaque study

#### Histopathology

Post-mortem examination of the extracted brain tissue did not reveal any macroscopic damage. A total of 12 H&E slides of brain tissue were evaluated: six slides, sampling left and right hemispheres, from two macaques. Histologic evaluation of tissue containing the focused ultrasound beam path from the four targeted locations did not show any evidence of damage in either macaque (Fig. S1). Specifically evaluated parameters included necrosis, apoptosis, edema, hemorrhage, inflammation, and neuropil rarefaction. Red blood cell extravasation could not be evaluated as these animals were perfused (*i.e.*, exsanguinated) prior to histologic examination.

### Sheep study

Estimates of *in situ* ultrasound intensity were obtained based on hydrophone measurements of pressure transmitted through each *ex vivo* skull cap. The acoustic power levels applied during the study corresponded to *in situ* peak pressure estimates of 1.7-3.6 MPa for MR-ARFI (Fig. 4(b)) and 0.25-0.9 MPa for neuromodulation (Fig. 4(e)), and *in situ* I_*SPTA*_ estimates ranging from 5.6-26.5 W/cm^2^ for MR-ARFI (Fig. 4(c)) and 0.6-13.8 W/cm^2^ for neuromodulation (Fig. 4(f)).

The number of FUS bursts applied to each location are stratified by the estimated *in situ* peak pressure and intensity of the sonication as shown in Fig. 5 for MR-ARFI and neuromodulation. Observations at multiple locations of the same number of bursts and estimated pressure or intensity are indicated by the color scale. High peak pressure values for MR-ARFI sonications were applied for short durations of 16 ms within the pulse repetition period, resulting in temporal average intensities that were similar to or slightly higher than the neuromodulation I_*SPTA*_ estimates, despite much lower neuromodulation peak pressures. In all sheep, transcranial FUS was confirmed by visualization of the focal spot by MR-ARFI with targeting to at least one subcortical location (Fig 6).

**Figure 5:**
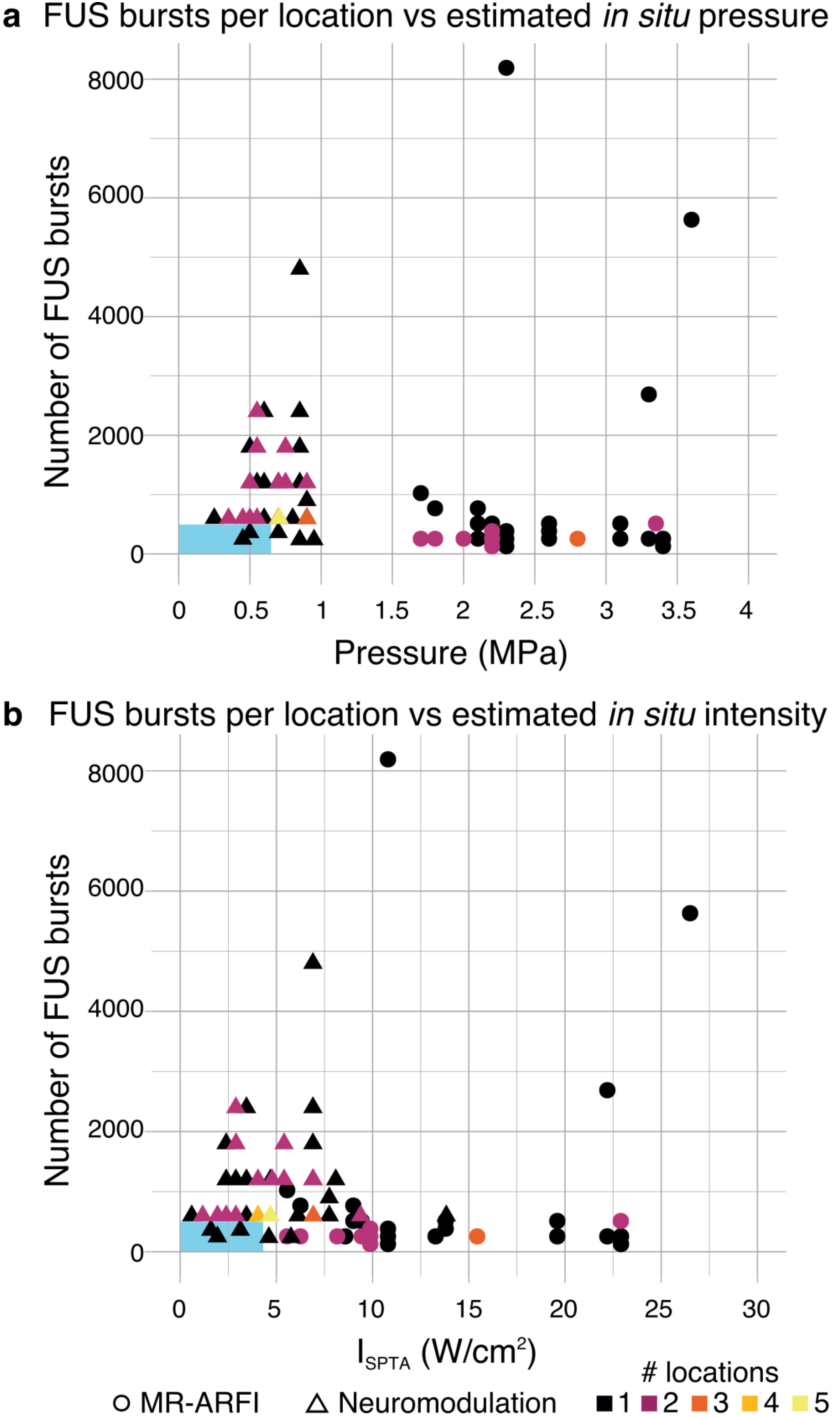
Distribution of the number of FUS bursts applied to each location with respect to the estimated *in situ* (a) peak pressure and (b) intensity of each sonication. MR-ARFI sonications (circles) were estimated to have *in situ* peak pressure between 1.7 and 3.6 MPa, which, due to the short 16 ms sonication times, corresponded to between 5.6 and 26.5 W/cm^2^ I_*SPTA*_. Neuromodulation sonications (triangles) were estimated to have peak *in situ* pressure between 0.25 and 0.9 MPa, corresponding to 0.6 and 13.8 W/cm^2^ I_*SPTA*_. The color scale indicates the number of locations at which each combination of *in situ* pressure or intensity and number of FUS bursts was observed. Blue rectangles indicate the range of parameters reported in human neuromodulation studies.

**Figure 6:**
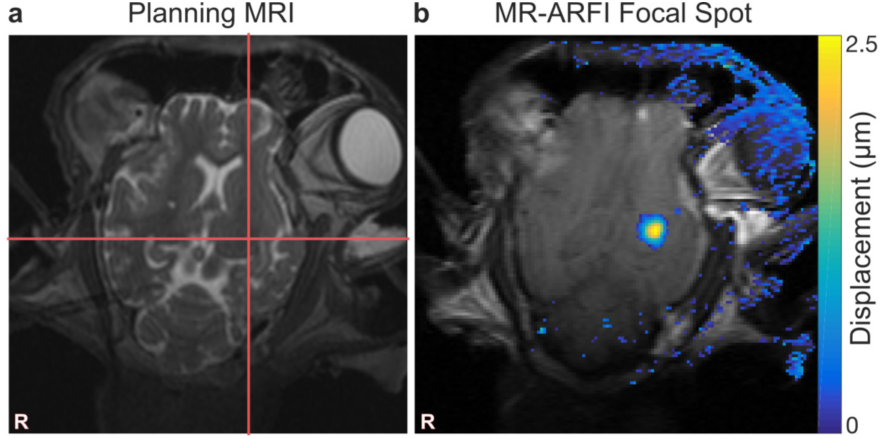
Focal spot targeting and visualization. (a) Prescribed focal spot is indicated by red cross hairs drawn on T2-weighted MRI. (b) Tissue displacement at the focal spot is shown as an overlay on the MR-ARFI magnitude image. Stray pixels in the displacement map outside the brain are artifact due to slight changes between two MR-ARFI acquisitions.

#### Histopathology

Overall, a total of 183 H&E slides of brain tissue from 13 sheep were evaluated for histologic damage. Of these, 128/183 received direct FUS exposure (sampled at the focal spot location and/or 3 mm rostral/caudal), 19/183 were internal controls (*i.e.*, contralateral hemisphere to that which received FUS), and 36/183 were experimental controls (*i.e.*, no FUS to either hemisphere). Overall, no FUS-related pre-mortem histologic findings were noted in any of the examined slides. Figure 7 summarizes the frequency of post-mortem histologic findings across study groups. The presence of each finding is reported for each hemisphere, where green boxes outline hemispheres that received FUS. The color scale represents the percentage of H&E slides that were positive for each histologic feature.

**Figure 7:**
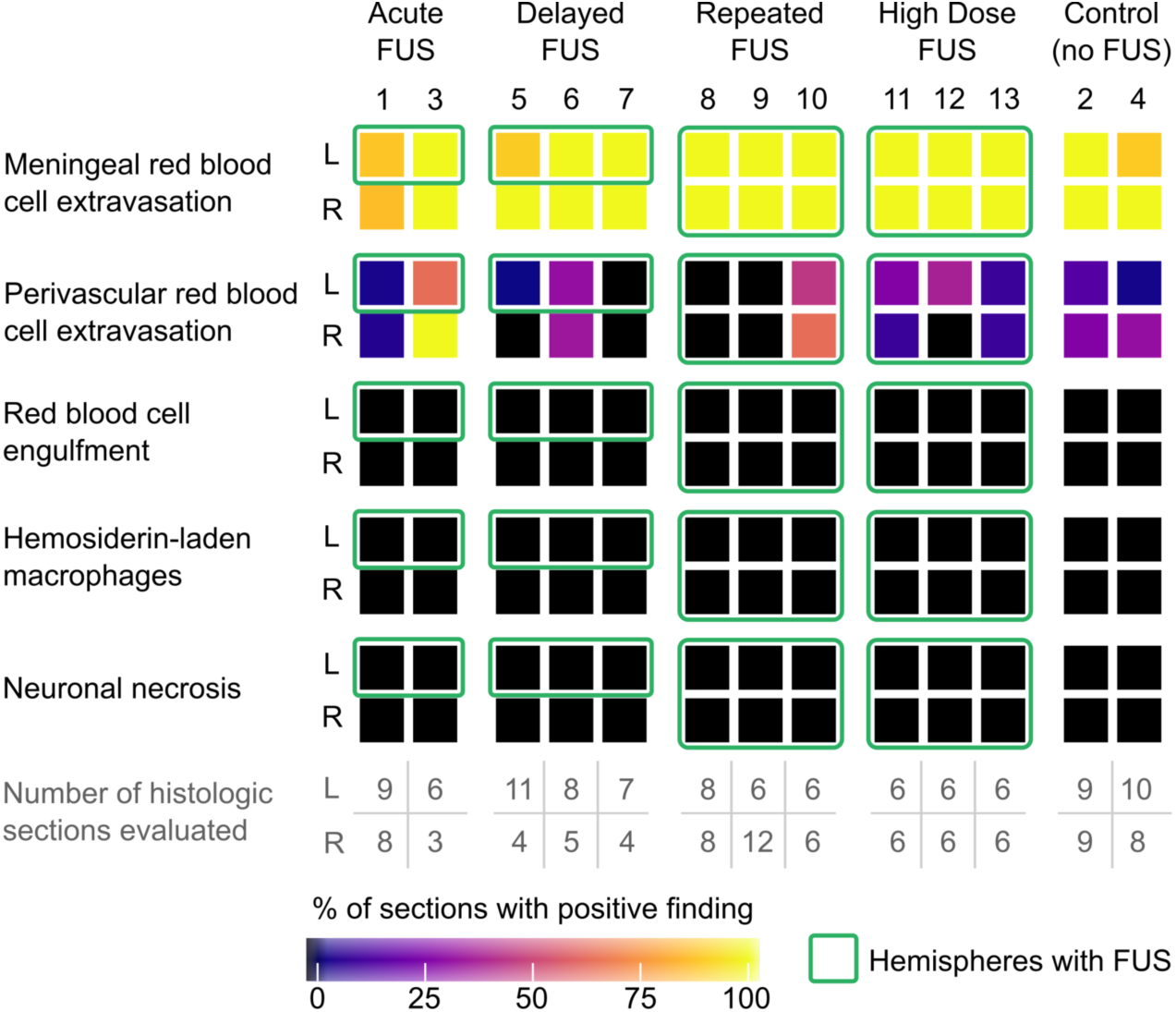
Prevalence of histologic findings within *in vivo* sheep study. The percentage of sections in which histologic findings were observed are reported for each animal by hemisphere (L and R; animal number listed at the top of each column). The number of histologic sections evaluated are reproduced from Fig. 2(a) for convenience. Green boxes indicate hemispheres where focused ultrasound was applied (all other boxes are internal controls or experimental controls). Meningeal and rare perivascular red blood cell extravasation were common histologic findings across all study groups, independent of whether any FUS was applied or which hemisphere was sonicated (in the case of FUS application). Necrosis, macrophage infiltration, red blood cell engulfment (erythrophagocytosis), and intracellular red blood cell breakdown (hemosiderin-laden macrophages), which would be expected to accompany true pre-mortem tissue damage, were not observed.

Histologic findings were limited to post-mortem red blood cell extravasation (meningeal or parenchymal) following brain extraction. Red blood cell extravasation was never observed at the precise sites of FUS targets. When present, parenchymal post-mortem red blood cell extravasations were randomly distributed within tissues distant to the FUS target. The number of incidences (foci) of scattered red blood cell extravasation in the parenchyma was quantified for each tissue section (Fig. 8). Our results suggest the rate of parenchymal red blood cell extravasation did not increase with FUS, but equivalence tests between FUS and control sections were not statistically significant. We performed a cluster-adjusted logistic regression and found the risk of red blood cell extravasation in the meninges is equivalent within +/-10% with p<0.05 between FUS treated and untreated tissue sections.

**Figure 8:**
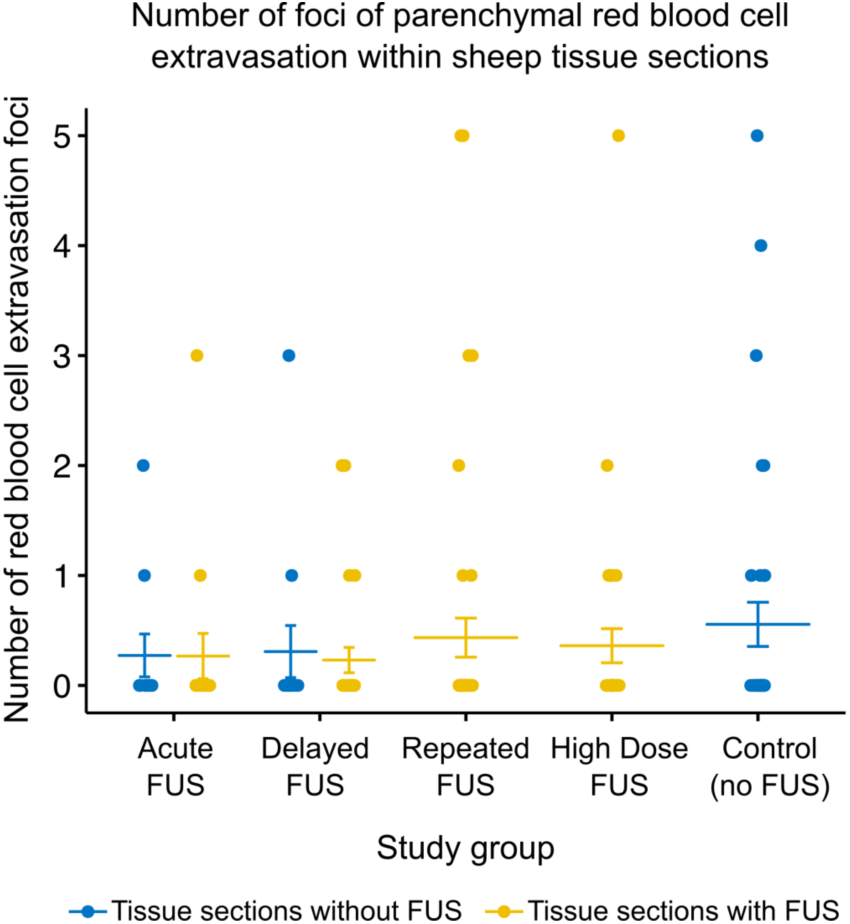
Summary of parenchymal red blood cell extravasation foci in H&E-stained sheep brain tissue slides. The number of foci per slide are shown for tissue taken from hemispheres without FUS (blue dots) and hemispheres with FUS (yellow dots) for each study group where applicable. Bars indicate mean and standard error.

#### Acute FUS group

Histologically, sheep euthanized less than 24 hours (n=2) following MRI and FUS exhibited red blood cell extravasation within the meninges (2/2) as well as rare perivascular red blood cells within neural parenchyma (2/2), regardless of hemispheric location (left vs right) and FUS application (Fig. 9(a,b,h,i)). No concurrent pre-mortem histologic findings (*i.e.*, necrosis, red blood cell engulfment (erythrophagocytosis), and intracellular red blood cell breakdown (hemosiderin-laden macrophages)) were noted in areas of red blood cell extravasation. However, acute hemorrhage can be histologically indistinguishable from post-mortem red blood cell extravasation (Finnie, 2016). Thus, a delayed euthanasia timepoint was established to confirm that red blood cell extravasation was indeed a post-mortem tissue extraction artifact rather than true pre-mortem hemorrhage.

**Figure 9:**
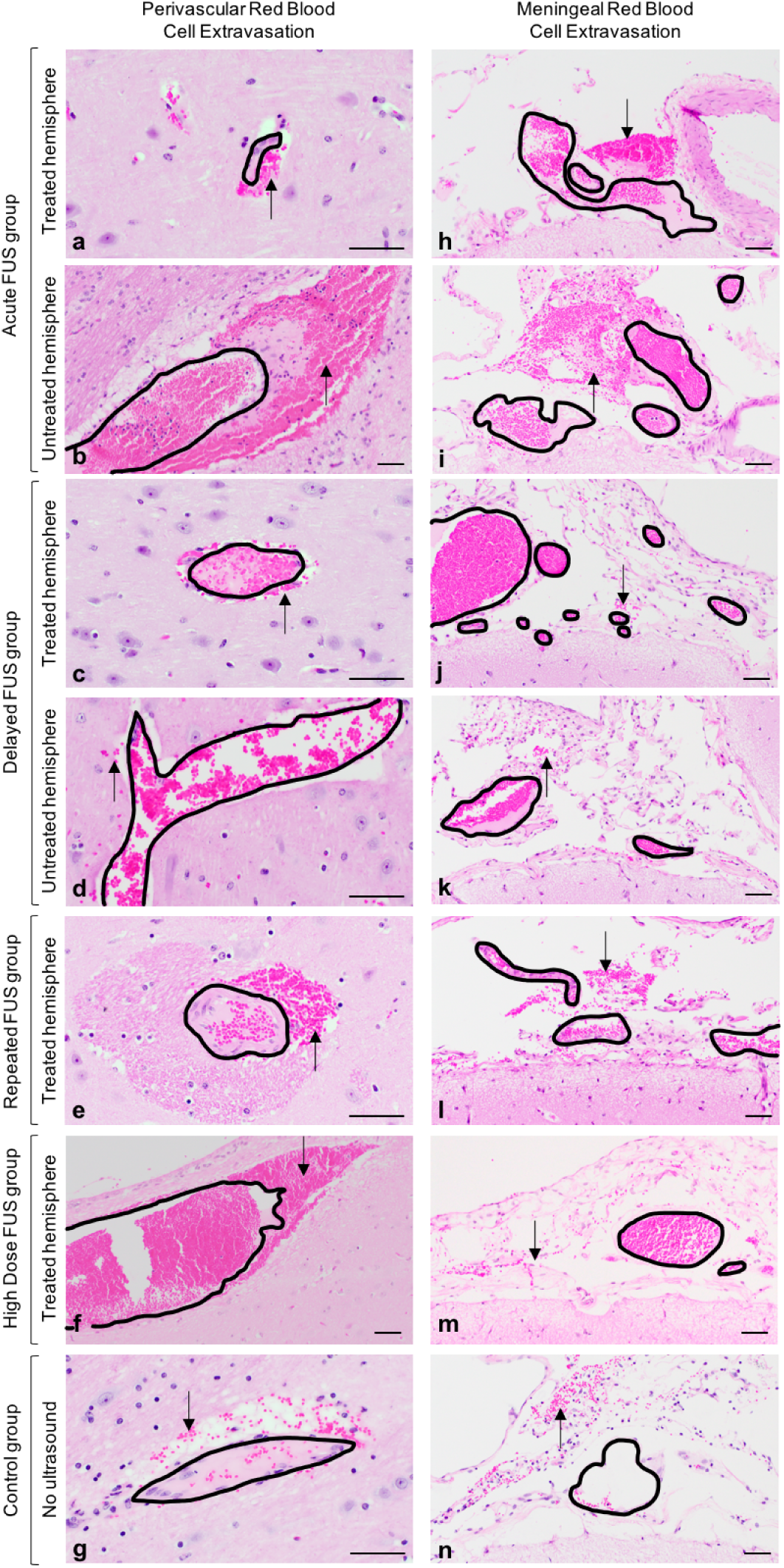
Post-mortem perivascular and meningeal red blood cell extravasation does not differ across sheep treatment groups. Randomly scattered small volumes of extravasated red blood cells (black arrows) were identified adjacent to blood vessels within the neural parenchyma (a-g) and throughout the meninges (h-n) regardless of ultrasound exposure. Black outlines indicate blood vessel walls and delineate intravascular from extravascular red blood cells. No red blood cell extravasation was observed at parenchymal locations targeted with FUS. No associated pre-mortem tissue reactions (*i.e.*, red blood cell engulfment (erythrophagocytosis), red blood cell breakdown (hemosiderosis), necrosis, or edema) were identified in any of the examined sections. Hematoxylin and eosin, scale bar = 50 *µ*m.

#### Delayed FUS group

In order to confirm that extravascular red blood cells seen in the acute FUS group reflected artifact following post-mortem tissue extraction, a delayed euthanasia timepoint was established (4- to 7-days post-FUS). In general, approximately 2- to 4-days following meningeal (or subarachnoid) hemorrhage, a normal response to hemorrhage should include erythrophagocytosis, while hemosiderin-laden macrophages are typically seen around 6- to 7-days post-hemorrhage (Finnie, 2016; Rao et al., 2016). In our study, sheep euthanized 96-168 following MRI and FUS exhibited extravascular red blood cells within the meninges (3/3) and rare extravascular red blood cells within neural parenchyma (2/3), regardless of hemispheric location (left vs right) and FUS application (Fig. 9(c,d,j,k)). Furthermore, at 96-168 hours following FUS, there was still no evidence of concurrent histologic abnormalities (such as those listed above) in regions of red blood cell extravasation.

#### Repeated FUS group

Tissue from sheep treated with FUS over multiple days exhibited extravascular red blood cells within the meninges (3/3) similar to the other groups. Occasional perivascular red blood cells were observed bilaterally within the neural parenchyma for one sheep (sheep 10; Fig. 9(e,l)). No other concurrent pre-mortem histologic findings (*i.e.*, necrosis, macrophage infiltration, red blood cell engulfment (erythrophagocytosis), and intracellular red blood cell breakdown (hemosiderin-laden macrophages)) were observed.

#### High dose FUS group

Sheep in the high dose group received prolonged consecutive MR-ARFI sonication to a single location on the last day of study, with the total number of MR-ARFI applications at the high dose location (25, 44, and 70 repetitions for sheep 11, 12, and 13, respectively) greatly exceeding the highest number of repetitions applied within the other FUS groups (8 repetitions for sheep 10). Neuromodulation sonications were similar to those applied in the other FUS groups. As with sheep in other groups, extravascular red blood cells were noted in the meninges (3/3) and rarely in parenchyma (3/3) (Fig. 9(f,m)). No other histologic findings accompanied extravascular red blood cells. Additionally, no histologic findings were observed at the high dose location or other locations targeted with FUS in any sheep. TUNEL results confirm no evidence of apoptosis at the high dose location for all three sheep (Fig. S2).

#### Control group

Control animals that only underwent the MRI procedure (*i.e.*, no FUS) also exhibited red blood cell extravasation within the meninges (2/2) and rarely within neural parenchyma (2/2) (Fig. 9(g,n)). As with sheep that underwent FUS, no evidence of concurrent premortem histologic findings (*i.e.*, necrosis, macrophage infiltration, red blood cell engulfment (erythrophagocytosis), and intracellular red blood cell breakdown (hemosiderinladen macrophages)) was observed in areas of red blood cell extravasation.

## Discussion

The results of this study suggest that the transcranial MR-ARFI and neuromodulation FUS protocols evaluated did not result in histologic tissue damage. No histologic abnormalities were observed at the site of FUS targets in either rhesus macaques or sheep, although post-mortem parenchymal red blood cell extravasation was observed in other brain regions of sheep tissue sections (*i.e.*, away from the focal spot).

Histologic findings were similar in both FUS treated and untreated hemispheres, as well as in control groups. Tissue sections from all sheep exhibited red blood cell extravasation in the meninges regardless of FUS application, treated hemisphere, or survival time (Fig 7). Through the process of post-mortem skull removal, meningeal blood vessels (*e.g.*, dural) are frequently ruptured resulting in the observed meningeal red blood cell extravasation. Furthermore, vibrations during extraction are strong enough to result in rare extravasations of red blood cells from parenchymal vessels. Multiple sections from both FUS (treated and untreated hemispheres) and control groups exhibited perivascular red blood cell extravasation in cortical tissue regions separate from those identified as FUS targets (Fig 8). No macrophage infiltration, erythrophagocytosis, hemosiderin-laden macrophages, tissue necrosis, or other indicators of tissue reactivity to damage were observed (Fig. 7), confirming post-mortem artifact.

Selecting appropriate euthanasia time points is crucial to interpreting histologic findings. At time points less than 24 hours, true small volume hemorrhage can be indistinguishable from tissue damage incurred during post-mortem brain extraction (Maxie, 2007). Following 72 hours, true pre-mortem hemorrhage should exhibit concurrent macrophage infiltration, erythrophagocytosis, and/or hemosiderin-laden macrophages (Rao et al., 2016). The absence of this expected tissue reactivity within our sheep cohort confirm that meningeal and extravascular red blood cells seen across both hemispheres and experimental groups were artifact due to post-mortem tissue extraction.

We evaluated *in situ* intensities similar to and slightly higher than previously reported I_*SPTA*_ values of up to 4.4 W/cm^2^ in humans, 9.5 W/cm^2^ in macaques, and 6.7 W/cm^2^ in sheep (Lee et al., 2016a; Verhagen et al., 2019; Lee et al., 2016c). The study in sheep reported microhemorrhage on H&E-stained tissue following 500 or more bursts of neuromodulation (300 ms long burst duration repeated in 1 second intervals at 50% duty cycle) at 3.3-5.7 W/cm^2^, but not at 6.7 W/cm^2^ I_*SPTA*_. Of fifteen publications assessing histology after neuromodulation, this was the only one to report abnormal findings, as summarized in a recent review of the ultrasound neuromodulation literature (Blackmore et al., 2019). However, because these foci of microhemorrhage were identified 4-64 days following treatment, with an absence of concurrent parenchymal reaction, we speculate that this finding may in fact be a post-mortem artifact.

In our study, repeated FUS neuromodulation and MR-ARFI sonications to the same focal spot location, either within one session or on multiple days, at various intensity levels, were not accompanied by histologic damage. We evaluated histology following a similar neuromodulation FUS protocol as Lee *et al.* In macaques, there was no tissue damage following 500 bursts at tissue locations receiving intensities of 0.4, 1.6, 6.4, and 25.8 W/cm^2^ I_*SPTA*_. Sonications of between 240 and 4800 bursts per location at intensity levels ranging from 0.6 and 13.8 W/cm^2^ I_*SPTA*_ did not result in pre-mortem damage in sheep. Furthermore, we evaluated histology from locations receiving between 128 and 8192 MR-ARFI bursts at a given intensity level, ranging from 5.6 and 26.5 W/cm^2^ I_*SPTA*_, and found no pre-mortem damage from either H&E- or TUNEL-stained tissue. One limitation of this study is that we did not detect tissue damage with either MR-ARFI or neuromodulation FUS.

Skull bone absorbs and dephases ultrasound which introduces a risk of cortical heating, and has been demonstrated to contribute to variations in FUS treatment across patients (Vyas et al., 2016). In our study, hydrophone measurements through *ex vivo* sheep skull caps resulted in a range of estimated *in situ* intensities, even when similar acoustic power levels were applied (Fig 4). Particular attention has been paid to thermal rise during neuromodulation, and a recent retrospective study has reported a simulated cortical temperature rise of 7°C caused by skull heating during preclinical neuromodulation (Constans et al., 2018). Several contemporary neuromodulation studies in humans have included assessments that no significant temperature rise in the brain is expected from skull heating with their protocols (Legon et al., 2014; Mueller et al., 2016; Ai et al., 2018; Legon et al., 2018a; Verhagen et al., 2019; Attali et al., 2019). We did not observe signs of cortical tissue damage due to skull heating in the rhesus macaque or sheep studies. Prior to treatment, simulations could be used to optimize FUS parameters to achieve a desired *in situ* intensity, and reduce the risk of tissue heating near bone (Mueller et al., 2016, 2017; Constans et al., 2018).

## Conclusions

The transcranial focused ultrasound protocols and equipment tested here did not result in pre-mortem tissue damage in rhesus macaques or sheep. Our study examined a range of experimental parameters including number of focal spot locations, number of FUS bursts applied to each spot, timing between FUS sessions, and applied acoustic intensity, exceeding the levels previously evaluated in other studies. Furthermore, we demonstrate that extravascular red blood cells may occur in extracted tissue whether or not focused ultrasound is applied. Results underscore the importance of selecting appropriate euthanasia timepoints and including experimental controls when interpreting histologic findings.

## Acknowledgements

The authors would like to thank Karla and Kevin Epperson for help with MRI, Benjamin Franco for help with sheep, Rachelle Bitton for help with MR-ARFI, Patrick Ye for help with hydrophone measurements, Jarrett Rosenberg for help with analyses, Megan Albertelli for help with rhesus macaques, Adrienne Mueller for rhesus macaque perfusions and tissue harvesting, Elias Godoy for sheep tissue harvesting, the Stanford Comparative Medicine Animal Histology Service Center for slide processing, and Insightec Ltd. for research support. This work was supported by NIH T32 EB009653, T32 CA009695, R01 MH111825, R01 EB019005, K99 NS100986.

---

**Figure S1:**
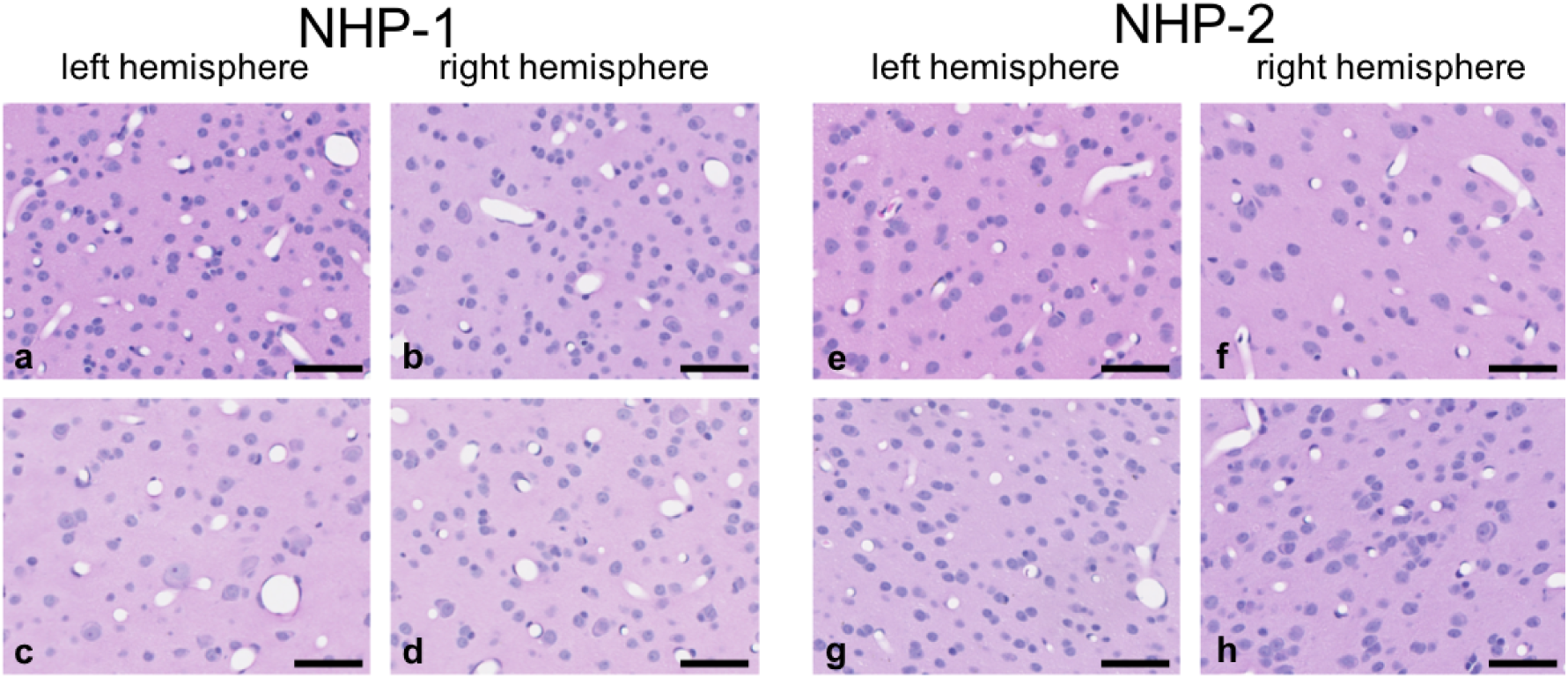
Histologic findings within *in vivo* NHP study. No histologic lesions were identified in NHP-1 (a-d) or NHP-2 (e-h). Representative normal cortical tissue is shown from FUS targeted regions corresponding to those shown in Figure 1. Hematoxylin and eosin, scale bar = 50 *µ*m.

**Figure S2:**
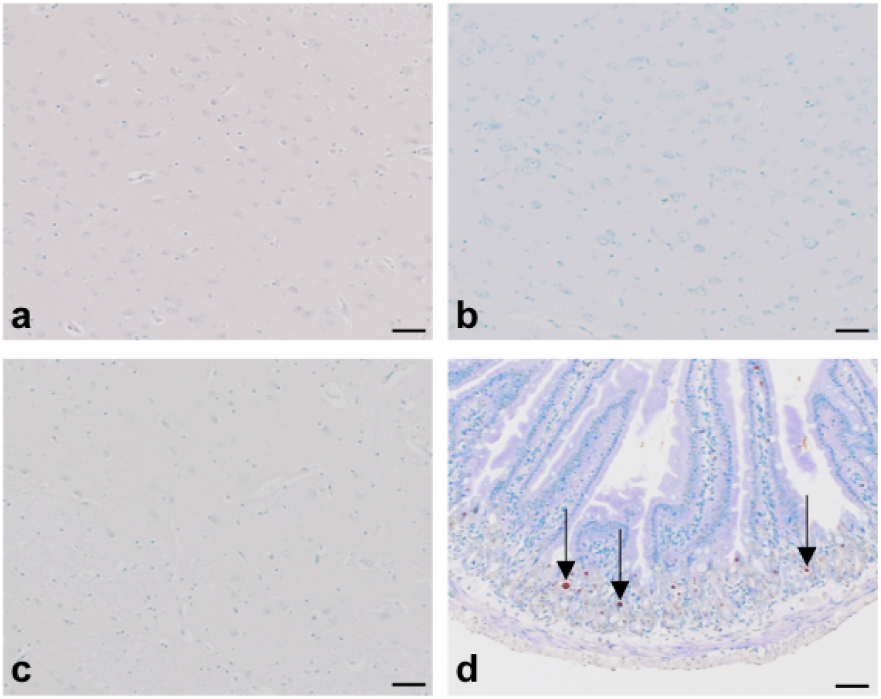
TUNEL staining in sheep in the high dose FUS group. Sheep 11 (a), 12 (b), and 13 (c) were TUNEL-negative at the targeted site of prolonged MR-ARFI repetitions. Figure d demonstrates apoptotic dark-brown, nuclear TUNEL-positivity (arrows) within the small intestinal epithelium of mice having undergone irradiation. TUNEL, scale bar = 50 *µ*m.

## References

Ai L, Bansal P, Mueller JK, Legon W. Effects of transcranial focused ultrasound on human primary motor cortex using 7T fMRI: a pilot study. BMC neuroscience, 2018;19:56.

Attali D, Houdouin A, Tanter M, Aubry JF. Thermal safety of transcranial focused simulation in human: retrospective numerical estimation of thermal rise in cortical and subcortical simulation setups. In: 19th International Symposium on Therapeutic Ultrasound, Barcelona, 2019.

Auboiroux V, Viallon M, Roland J, Hyacinthe JN, Petrusca L, Morel DR, Goget T, Terraz S, Gross P, Becker CD, et al. ARFI-prepared MRgHIFU in liver: Simultaneous mapping of ARFI-displacement and temperature elevation, using a fast GRE-EPI sequence. Magnetic resonance in medicine, 2012;68:932–946.

Bitton RR, Kaye E, Dirbas FM, Daniel BL, Pauly KB. Toward MR-guided high intensity focused ultrasound for presurgical localization: Focused ultrasound lesions in cadaveric breast tissue. Journal of magnetic resonance imaging, 2012;35:1089–1097.

Blackmore J, Shrivastava S, Sallet J, Butler CR, Cleveland RO. Ultrasound neuromodulation: A review of results, mechanisms and safety. Ultrasound in medicine & biology, 2019.

Chaplin V, Phipps M, Jonathan S, Grissom W, Yang P, Chen L, Caskey C. On the accuracy of optically tracked transducers for image-guided transcranial ultrasound. International journal of computer assisted radiology and surgery, 2019:1–11.

Constans C, Mateo P, Tanter M, Aubry JF. Potential impact of thermal effects during ultrasonic neurostimulation: retrospective numerical estimation of temperature elevation in seven rodent setups. Physics in Medicine & Biology, 2018;63:025003.

Dallapiazza RF, Timbie KF, Holmberg S, Gatesman J, Lopes MB, Price RJ, Miller GW, Elias WJ. Noninvasive neuromodulation and thalamic mapping with low-intensity focused ultrasound. Journal of neurosurgery, 2017:1–10.

Deffieux T, Younan Y, Wattiez N, Tanter M, Pouget P, Aubry JF. Low-intensity focused ultrasound modulates monkey visuomotor behavior. Current Biology, 2013;23:2430–2433.

Finnie JW. Forensic pathology of traumatic brain injury. Veterinary pathology, 2016;53:962–978.

Fomenko A, Neudorfer C, Dallapiazza RF, Kalia SK, Lozano AM. Low-intensity ultrasound neuromodulation: an overview of mechanisms and emerging human applications. Brain stimulation, 2018.

Hameroff S, Trakas M, Duffield C, Annabi E, Gerace MB, Boyle P, Lucas A, Amos Q, Buadu A, Badal JJ. Transcranial ultrasound (tus) effects on mental states: a pilot study. Brain stimulation, 2013;6:409–415.

Holbrook AB, Ghanouni P, Santos JM, Medan Y, Butts Pauly K. In vivo mr acoustic radiation force imaging in the porcine liver. Medical physics, 2011;38:5081–5089.

Holbrook AB, Santos JM, Kaye E, Rieke V, Pauly KB. Real-time MR thermometry for monitoring HIFU ablations of the liver. Magnetic Resonance in Medicine, 2010;63:365–373.

Huang Y, Curiel L, Kukic A, Plewes DB, Chopra R, Hynynen K. Mr acoustic radiation force imaging: in vivo comparison to ultrasound motion tracking. Medical physics, 2009;36:2016–2020.

Kubanek J. Neuromodulation with transcranial focused ultrasound. Neurosurgical focus, 2018;44:E14.

Larrat B, Pernot M, Aubry JF, Dervishi E, Sinkus R, Seilhean D, Marie Y, Boch AL, Fink M, Tanter M. MR-guided transcranial brain HIFU in small animal models. Physics in medicine & biology, 2009;55:365.

Lee W, Chung YA, Jung Y, Song IU, Yoo SS. Simultaneous acoustic stimulation of human primary and secondary somatosensory cortices using transcranial focused ultrasound. BMC neuroscience, 2016a;17:68.

Lee W, Kim H, Jung Y, Song IU, Chung YA, Yoo SS. Image-guided transcranial focused ultrasound stimulates human primary somatosensory cortex. Scientific reports, 2015;5:8743.

Lee W, Kim HC, Jung Y, Chung YA, Song IU, Lee JH, Yoo SS. Transcranial focused ultrasound stimulation of human primary visual cortex. Scientific reports, 2016b;6:34026.

Lee W, Lee SD, Park MY, Foley L, Purcell-Estabrook E, Kim H, Fischer K, Maeng LS, Yoo SS. Image-guided focused ultrasound-mediated regional brain stimulation in sheep. Ultrasound in Medicine and Biology, 2016c;42:459–470.

Legon W, Ai L, Bansal P, Mueller JK. Neuromodulation with single-element transcranial focused ultrasound in human thalamus. Human brain mapping, 2018a;39:1995–2006.

Legon W, Bansal P, Tyshynsky R, Ai L, Mueller JK. Transcranial focused ultrasound neuromodulation of the human primary motor cortex. Scientific Reports, 2018b;8:10007.

Legon W, Sato TF, Opitz A, Mueller J, Barbour A, Williams A, Tyler WJ. Transcranial focused ultrasound modulates the activity of primary somatosensory cortex in humans. Nature neuroscience, 2014;17:322.

Lipsman N, Mainprize TG, Schwartz ML, Hynynen K, Lozano AM. Intracranial applications of magnetic resonance-guided focused ultrasound. Neurotherapeutics, 2014;11:593–605.

Marsac L, Chauvet D, Larrat B, Pernot M, Robert B, Fink M, Boch AL, Aubry JF, Tanter M. Mr-guided adaptive focusing of therapeutic ultrasound beams in the human head. Medical physics, 2012;39:1141–1149.

Martin E, Werner B. Focused ultrasound surgery of the brain. Current Radiology Reports, 2013;1:126–135.

Maxie MG. Jubb, Kennedy & Palmer’s pathology of domestic animals. 5th Edition. Elsevier, Edinburgh, 2007. p. 300.

McDannold N, Maier SE. Magnetic resonance acoustic radiation force imaging. Medical physics, 2008;35:3748–3758.

Mohammadjavadi M, Gaur P, Kubanek J, Popelka G, Pauly KB. Transcranial focused ultrasound neuromodulation of the visual system in a large animal (sheep). In: 19th International Symposium on Therapeutic Ultrasound, Barcelona, 2019.

Monti MM, Schnakers C, Korb AS, Bystritsky A, Vespa PM. Non-invasive ultrasonic thalamic stimulation in disorders of consciousness after severe brain injury: a first-in-man report. Brain Stimul, 2016;9:940–941.

Mueller JK, Ai L, Bansal P, Legon W. Computational exploration of wave propagation and heating from transcranial focused ultrasound for neuromodulation. Journal of neural engineering, 2016;13:056002.

Mueller JK, Ai L, Bansal P, Legon W. Numerical evaluation of the skull for human neuromodulation with transcranial focused ultrasound. Journal of neural engineering, 2017;14:066012.

Naor O, Krupa S, Shoham S. Ultrasonic neuromodulation. Journal of neural engineering, 2016;13:031003.

Rao MG, Singh D, Vashista RK, Sharma SK. Dating of acute and subacute subdural haemorrhage: a histo-pathological study. Journal of clinical and diagnostic research: JCDR, 2016;10:HC01.

Tyler WJ, Lani SW, Hwang GM. Ultrasonic modulation of neural circuit activity. Current opinion in neurobiology, 2018;50:222–231.

Verhagen L, Gallea C, Folloni D, Constans C, Jensen DE, Ahnine H, Roumazeilles L, Santin M, Ahmed B, Lehericy S, et al. Offline impact of transcranial focused ultrasound on cortical activation in primates. Elife, 2019;8:e40541.

Vyas U, Ghanouni P, Halpern CH, Elias J, Pauly KB. Predicting variation in subject thermal response during transcranial magnetic resonance guided focused ultrasound surgery: Comparison in seventeen subject datasets. Medical physics, 2016;43:5170–5180.

Vyas U, Kaye E, Pauly KB. Transcranial phase aberration correction using beam simulations and mr-arfi. Medical physics, 2014;41.

